# Cheminformatics Analysis of Natural Product Scaffolds: Comparison of Scaffolds Produced by Animals, Plants, Fungi and Bacteria

**DOI:** 10.1101/2020.01.28.922955

**Authors:** Peter Ertl, Tim Schuhmann

**Affiliations:** Novartis Institutes for BioMedical Reseach, Novartis Campus, CH-4056, Basel, Switzerland

**Keywords:** natural products, secondary metabolites, scaffolds, substructure analysis

## Abstract

Natural products (NPs) have evolved over a very long natural selection process to form optimal interactions with biologically relevant macromolecules. NPs are therefore an extremely useful source of inspiration for the design of new drugs. In the present study we report the results of a cheminformatics analysis of a large database of NP structures focusing on their scaffolds. First, general differences between NP scaffolds and scaffolds from synthetic molecules are discussed, followed by a comparison of the properties of scaffolds produced by different types of organisms. Scaffolds produced by plants are the most complex and those produced by bacteria differ in many structural features from scaffolds produced by other organisms. The results presented here may be used as a guidance in selection of scaffolds for the design of novel NP-like bioactive structures or NP-inspired libraries.

## 1 Introduction

Natural products (NPs) have been optimized in a very long natural selection process for optimal interactions with biologically relevant macromolecules. NPs are therefore an excellent source of substructures for the design of new drugs. Indeed, many drugs in the current pharmacopeia are NPs and many others are of NP origin (Newman and Cragg, 2016). In recent years we can witness a real explosion of interest in the use of NPs in drug discovery (Rodrigues *et al.*, 2016), (Harvey, Edrada-Ebel and Quinn, 2015).This may partly be attributed to the fact that high expectations in several novel technologies that have been introduced into the drug discovery process a decade ago have not fully materialized. These technologies, including combinatorial chemistry, high throughput screening and various –omics techniques, although improving the efficiency of the drug discovery process, did not fill development pipelines of pharmaceutical companies with a flood of new drug candidates as originally expected.

Another area where the NP scaffolds serve as inspiration for drug discovery is the design of small focused NP-like libraries (Davison and Brimble, 2019), (Grabowski, Baringhaus and Schneider, 2008). It is well known that the first generation of combinatorial libraries, containing mostly large, hydrophobic molecules with many rotatable bonds, was rather a disappointment concerning their biological activity. As a conclusion, chemists realized that not only the number of molecules synthesized is important, but also their properties and scaffold diversity.(Sauer and Schwarz, 2003) This led to re-evaluation of combichem design strategies, introduction of so called “diversity oriented synthesis” (DOS) (Tan, 2005) and “biology-oriented synthesis” (BIOS) (Wilk *et al.*, 2010). These methods aim to generate libraries of diverse small molecules to explore untapped and underrepresented regions of chemical space and inspiration by NP structures plays an important role in both these techniques.

The goal of the present study is to analyze the scaffolds present in NP molecules, compare them with scaffolds from synthetic molecules and identify typical structural characteristics of scaffolds present in metabolites of various organism classes. A comparison of the physicochemical properties of scaffolds was not included in this analysis, since this topic is already covered by several publications. (Wetzel *et al.*, 2007), (Ertl and Schuffenhauer, 2008).

## 2 Methodology

### 2.1 Data

The main goal of our study was to analyze scaffolds present in molecules produced by living organisms. As a data source we used natural product structures from the CRC Dictionary of Natural Products (DNP) (‘Dictionary of natural products 27.1, CRC Press, Taylor & Francis Group, Boca Raton, FL, USA’, no date). Since we wanted to perform the analysis with respect to the class of organisms producing the particular molecules only those database entries where the producing organism could be identified were retained. These were identified with the help of the Taxonomy Database of the National Center for Biotechnology Information.(*Taxonomy - NCBI*, no date) A Python script was used to analyze information from the biological source field (BSRC) of the molecules to identify the scientific name of the producing organism and to match it to the entry in the taxonomy database. This allowed assigning the source organism to one of the four classes (animals, plants, fungi or bacteria). If no scientific name could be found the Python script examined the words given in the source description field and compared them with the keywords typical for the various origin classes. For example “extracted from leaves” identified the origin as a plant, or “isolated from marine sponge” as an animal. This procedure allowed us to identify 129,794 NPs of plant origin, 20,842 of animal origin, 18,264 NPs produced by fungi and 13,201 by bacteria. This distribution reflects the fact that for a long time the main source of natural products were plant extracts. Later products from bacteria and fungi, mainly thanks to their antibiotic activity, gained popularity. Currently a very promising and fast growing field is the isolation of NPs from marine invertebrates (Carroll *et al.*, 2019). One reason for this is the enormous diversity of marine ecosystems, which provides a source of a large number of novel, diverse NP structures. Another reason might be the fact that marine NPs when used as “chemical warfare” agents by their producers dilute rapidly in water, and therefore they need to be very potent. However, one needs to be aware that the class “animals” used in the present analysis is highly heterogeneous and includes species ranging from marine invertebrates to mammals and thus results need to be interpreted keeping this in mind.

### 2.1 Molecule Processing

The molecular structures for which the producing organism could be identified were converted into SMILES format, cleaned by normalizing charges and by removing smaller parts (counterions, etc.). Before the actual scaffold analysis could be performed an additional processing step was necessary, namely in silico deglycosylation, i.e. removal of sugar units from the molecules. One major role of sugar moieties in NPs is to change the pharmacokinetic properties of the parent structures and make them more soluble (Elshahawi *et al.*, 2015). In most cases the sugar units do not directly influence the biological activity of the aglycon (although several important exceptions of this general rule exist). The sugar units are one of the most typical substructures present in NP molecules and since we were interested in the analysis of aglycon scaffolds the sugar units had to be removed before the actual scaffold extraction. The deglycosylation process we used is fully described in one of our previous studies (Koch *et al.*, 2005). In short, a recursive deglycosylation procedure that removed sugar units was applied, starting at the molecule periphery and iteratively progressing until no more sugars were present. From the resulted aglycons the scaffolds were extracted for the actual analysis. The term “scaffold” is used in medicinal chemistry literature rather freely, lacking clear definition. The exact meaning of this term varies from publication to publication and depends also on the particular area of interest. Throughout this article the term “scaffold” is used to describe part of the molecule that remains after removal of sugars and non-ring substituents, keeping, however exocyclic and exochain multiple bonds. This is the same definition that was applied to analyze scaffolds in our earlier studies (Ertl, 2014), (Ertl and Schuffenhauer, 2008).

The scaffolds extracted from NPs were compared with those of standard synthetic molecules represented by over 10 million drug-like commercially available samples from the ZINC database (Sterling and Irwin, 2015). These molecules were processed in the same way as the NP structures.

## 3 Results and Discussion

### 3.1 Difference Between Natural and Synthetic Scaffolds

In the first part of the study differences between natural and synthetic scaffolds were analysed. The 50 most frequent scaffolds from both sets (91 unique scaffolds totally) are shown in Figure 1. The horizontal axis in the diagram represents scaffold frequency (the most common scaffolds are on the left, less common ones on the right), whereas the vertical axis indicates the propensity of scaffolds being of natural (green area at the top) or synthetic (red area at the bottom) origin. This type of visualization nicely shows the differences between both sets. Scaffolds typical for NPs contain mostly aliphatic rings and relatively few heteroatoms with the most frequent one being oxygen. The most common structural features of synthetic scaffolds are phenyl rings, amide and sulfonamide linkers. They contain also much more heteroatoms, including oxygen, nitrogen as well as sulfur. Out of the 50 most common NP scaffolds 26 contain oxygen atoms, 5 nitrogen atoms and none sulfur atoms leaving 20 hydrocarbons scaffolds without any heteroatoms. In the synthetic scaffold set 40 scaffolds contain nitrogen, 32 oxygen and 11 sulfur atoms leaving only 3 hydrocarbon scaffolds. Out of the 50 most common synthetic scaffolds 47 contain aromatic rings, about two times as much as the NP scaffolds (26). Although this simple comparison was made only for the 50 most common scaffolds from each set, it reflects quite well the general differences between the scaffolds originated from NPs and synthetic molecules, respectively. A more detailed comparison of structural feature using larger sets of scaffolds can be found in Table 1.

**Table 1.** Selected substructure parameters for the 1000 most common scaffolds derived from animal (A), plant (P), fungal (F) and bacterial (B) metabolites as well as synthetic molecules (SM).

| Structural feature | A | P | F | B | SM |
| --- | --- | --- | --- | --- | --- |
| average number of atoms in scaffold | 19.7 | 18.3 | 18.3 | 25.6 | 17.0 |
| average number of rings in scaffold | 3.0 | 3.7 | 3.3 | 3.2 | 2.4 |
| average number of ring systems in scaffold | 1.2 | 1.4 | 1.3 | 1.6 | 2.2 |
| average number of stereocenters in scaffold | 2.5 | 2.9 | 2.0 | 1.6 | 0.1 |
| % of scaffolds with 7 or 8 membered rings | 8.6 | 10.1 | 7.7 | 3.2 | 1.0 |
| % of heteroatoms in scaffolds | 12.9 | 11.2 | 16.3 | 21.3 | 21.4 |
| % scaffolds containing aromatic ring | 32.5 | 49.9 | 57.2 | 63.1 | 98.1 |
| % scaffolds containing macrocycles | 22.5 | 6.4 | 12.2 | 34.3 | 0.0 |
| % scaffolds containing spiro system | 12.6 | 18.9 | 17.2 | 8.6 | 0.2 |
| % scaffolds being hydrocarbons | 16.2 | 16.3 | 8.9 | 4.0 | 2.0 |
| % scaffolds containing oxygen | 73.2 | 75.8 | 88.3 | 87.4 | 80.7 |
| % scaffolds containing nitrogen | 29.6 | 17.9 | 27.3 | 59.8 | 92.9 |
| % scaffolds containing sulfur | 2.5 | 0.3 | 2.0 | 7.5 | 30.0 |

**Figure 1.**
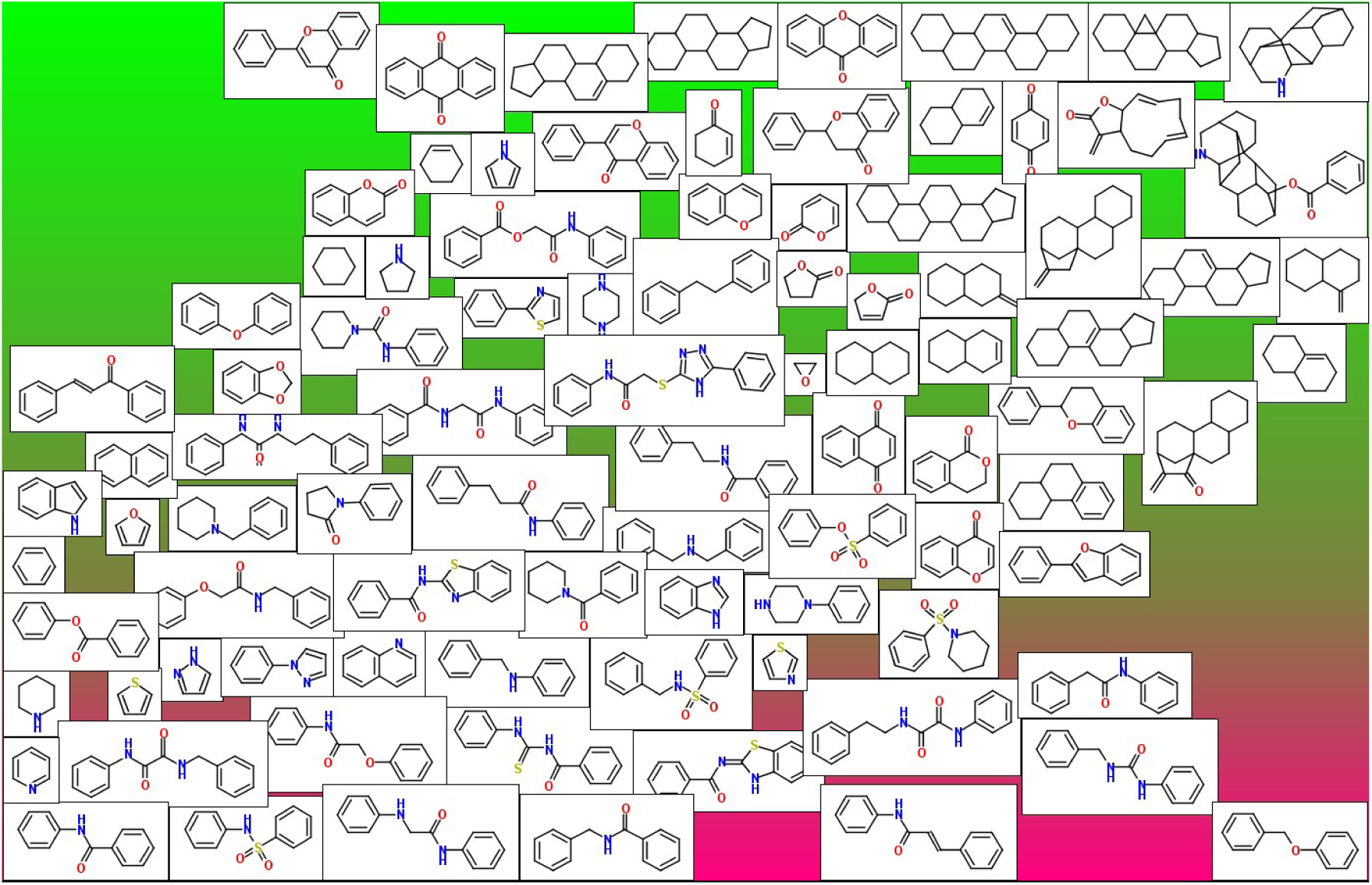
Plot of common scaffolds displaying their preference for natural products (green area) and synthetic molecules (red area). Position on the horizontal axis is proportional to the frequency of scaffolds - the most common scaffolds are on the left, less common ones on the right.

### 3.2 Differences Between Scaffolds Produced by Different Organism Classes

In the second part of the study differences between scaffolds from molecules produced by different classes of organisms were analysed. In Figures 2 - 5 the most typical scaffolds produced by different organisms are shown. These were obtained by ranking scaffolds in each class by their frequency and then removing those that were amongst the top 1000 also in at least 2 other classes. In this way the “promiscuous” NP scaffolds (including common coumarins, flavones, chromones, chromanes, quinones, steroids etc.) were removed. Although in the figures only 30 representative examples for each class are shown, a visual comparison even of this relatively small number of structures already indicates interesting differences. To better quantify the differences between producer organisms several simple substructure descriptors for the 1000 most frequent scaffolds from each class were calculated. They are listed together with the same descriptors for synthetic scaffolds in Table 1. Several interesting conclusions can be made based on the values in this table. The metabolites produced by bacteria contain the largest scaffolds (average number of atoms in bacterial scaffolds is 26 compared to 17-19 for other classes). This is mostly caused by the presence of macrocycles in a substantial portion (34.3%) of bacterial scaffolds, much more than in other scaffold classes (animal scaffolds 22.4%, fungal scaffolds 12.2% and plant scaffolds 6.4% only). Bacterial scaffolds also contain the largest portion of heteroatoms (21.3%). Out of the 1000 most frequent bacterial scaffolds 59.8% contain nitrogen (considerably more compared with the second largest portion 29.6% in animal metabolites) and are the only class where larger percentage of scaffolds contain sulfur (7.5%). In contrast to this, plant metabolites contain the smallest portion of heteroatoms (only 11.2%). The plant scaffolds are sterically most complex, containing 18.9% spirocyclic systems, followed closely by scaffolds produced by fungi (17.2%) and they contain also the largest proportion of flexible 7- and 8-membered rings (10.1%). Also with respect to the presence of stereocenters the plant scaffolds are the most complex (on average 2.9 stereo centers per scaffold), followed by animal (2.5), fungal (2.0) and bacterial (1.6) scaffolds, differing clearly from synthetic scaffolds that contain on average only 0.1 stereo centers per scaffold.

**Figure 2.**
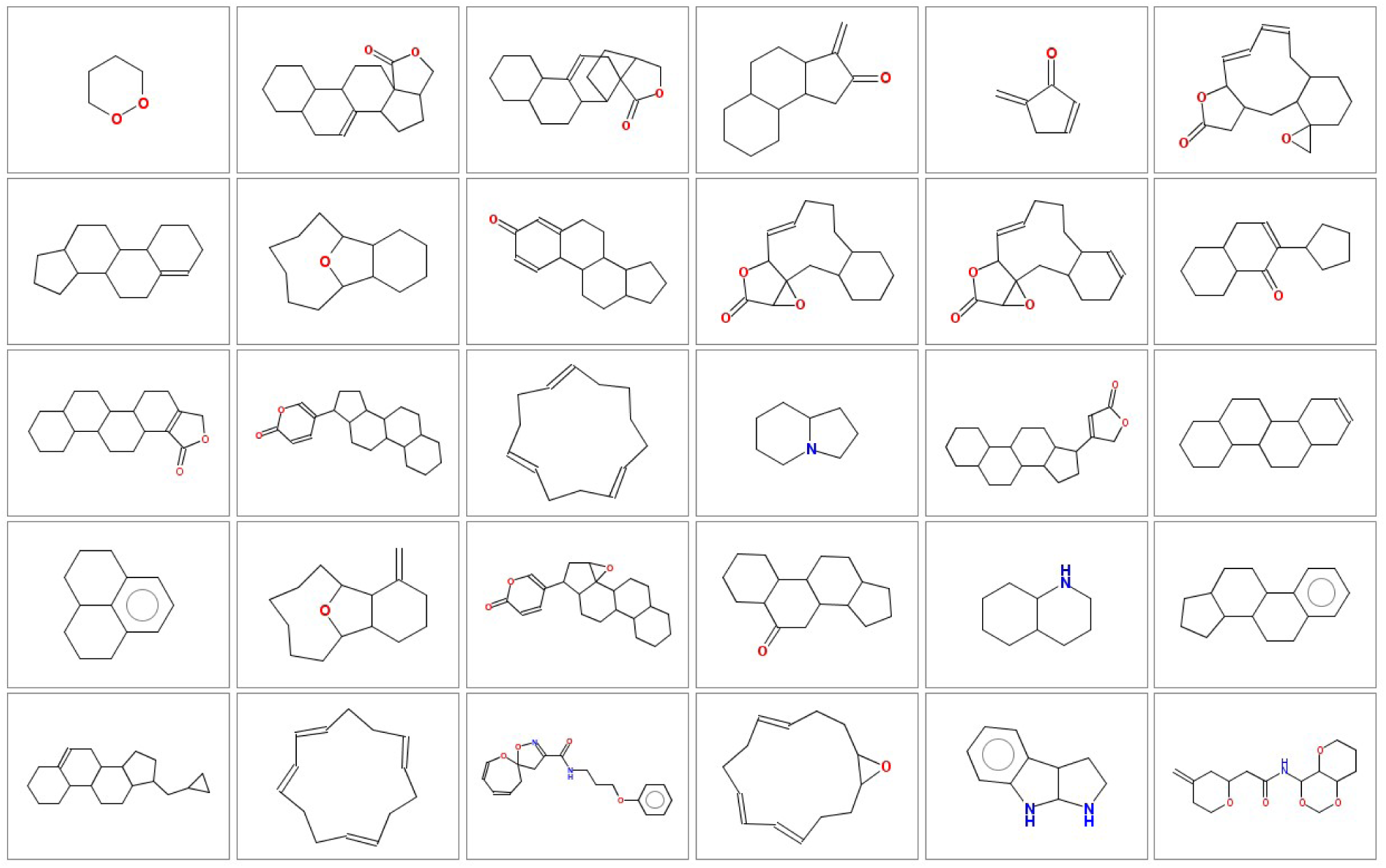
Typical scaffolds from molecules produced by animals.

**Figure 3.**
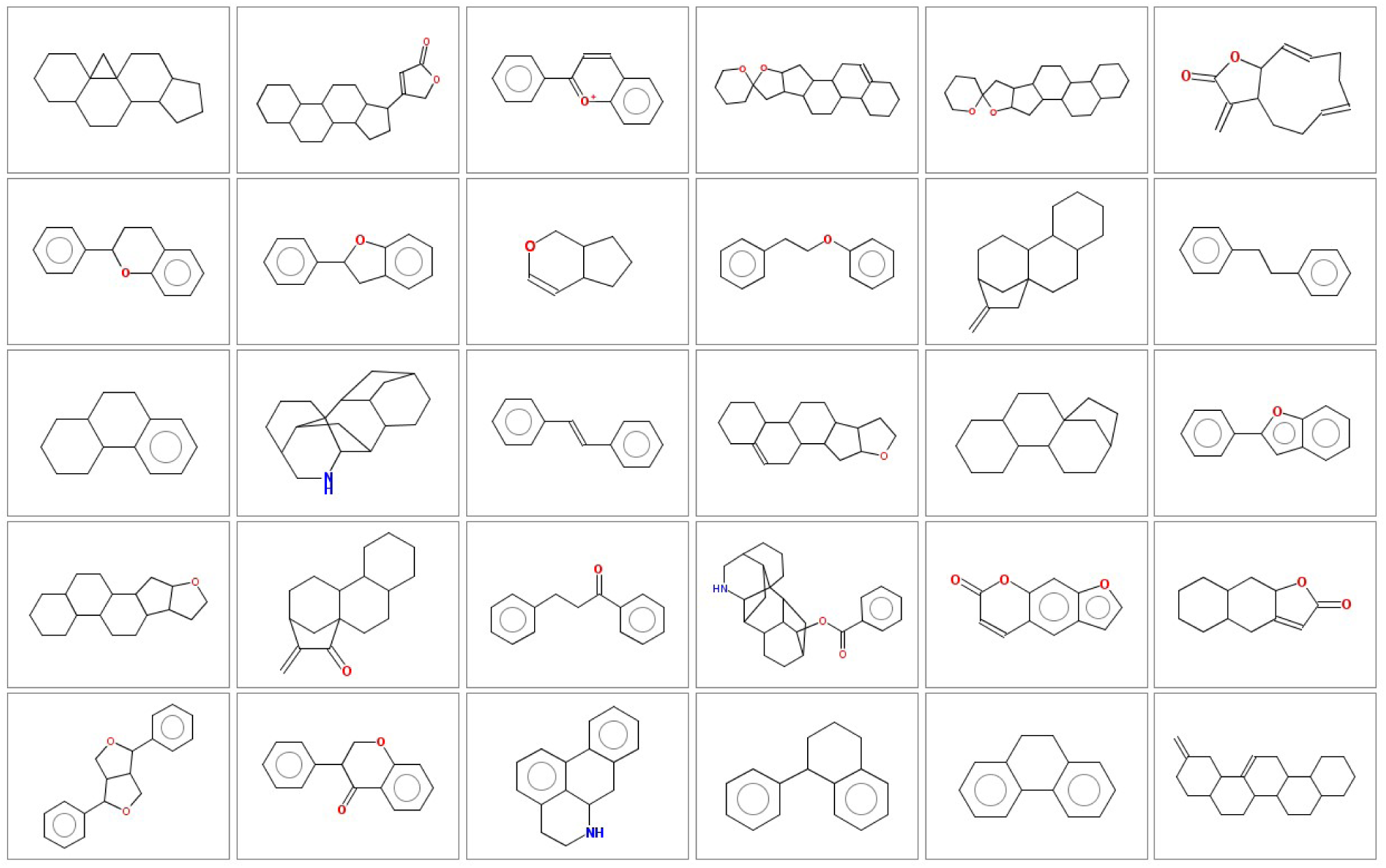
Typical scaffolds from molecules produced by plants.

**Figure 4.**
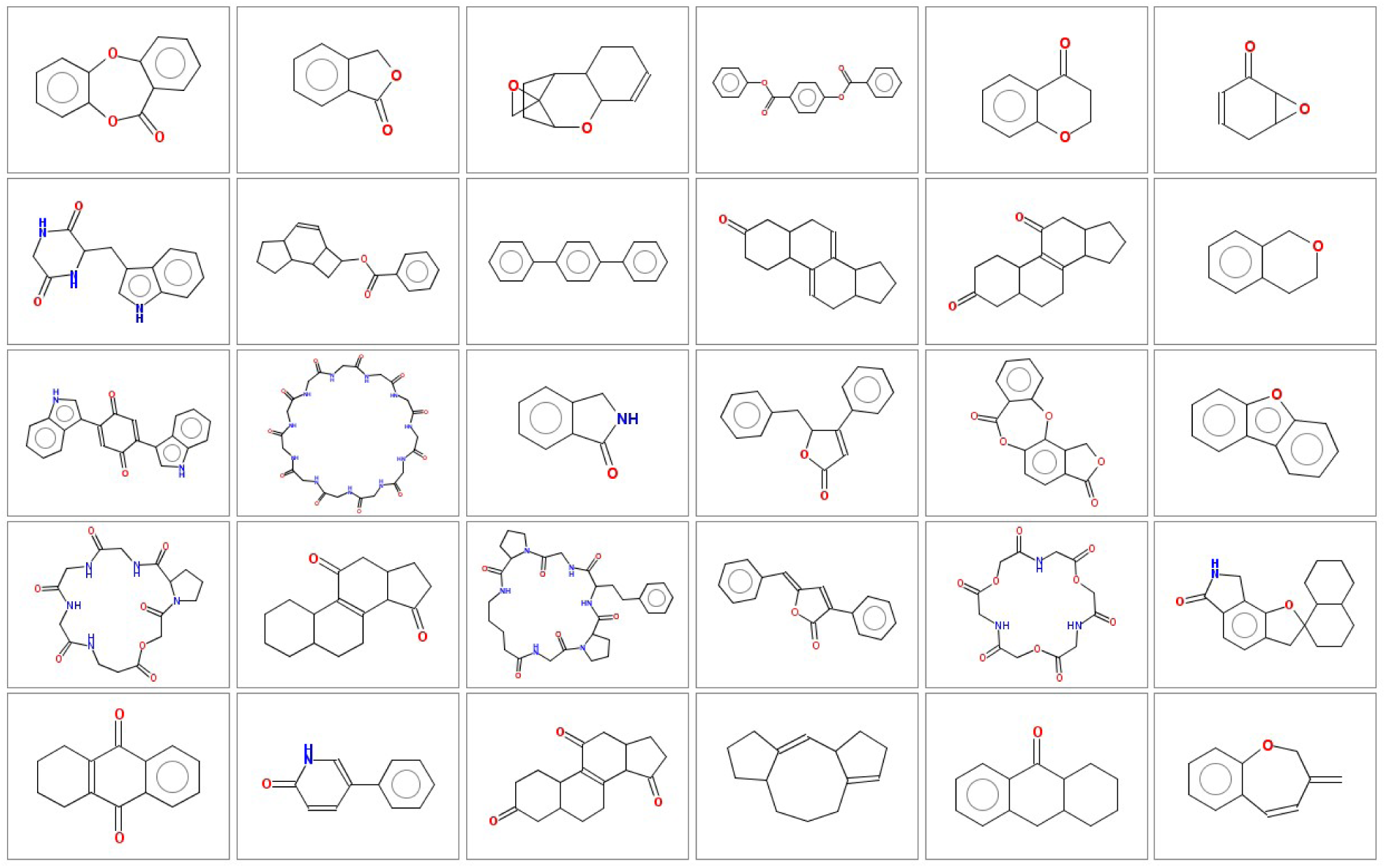
Typical scaffolds from molecules produced by fungi.

**Figure 5.**
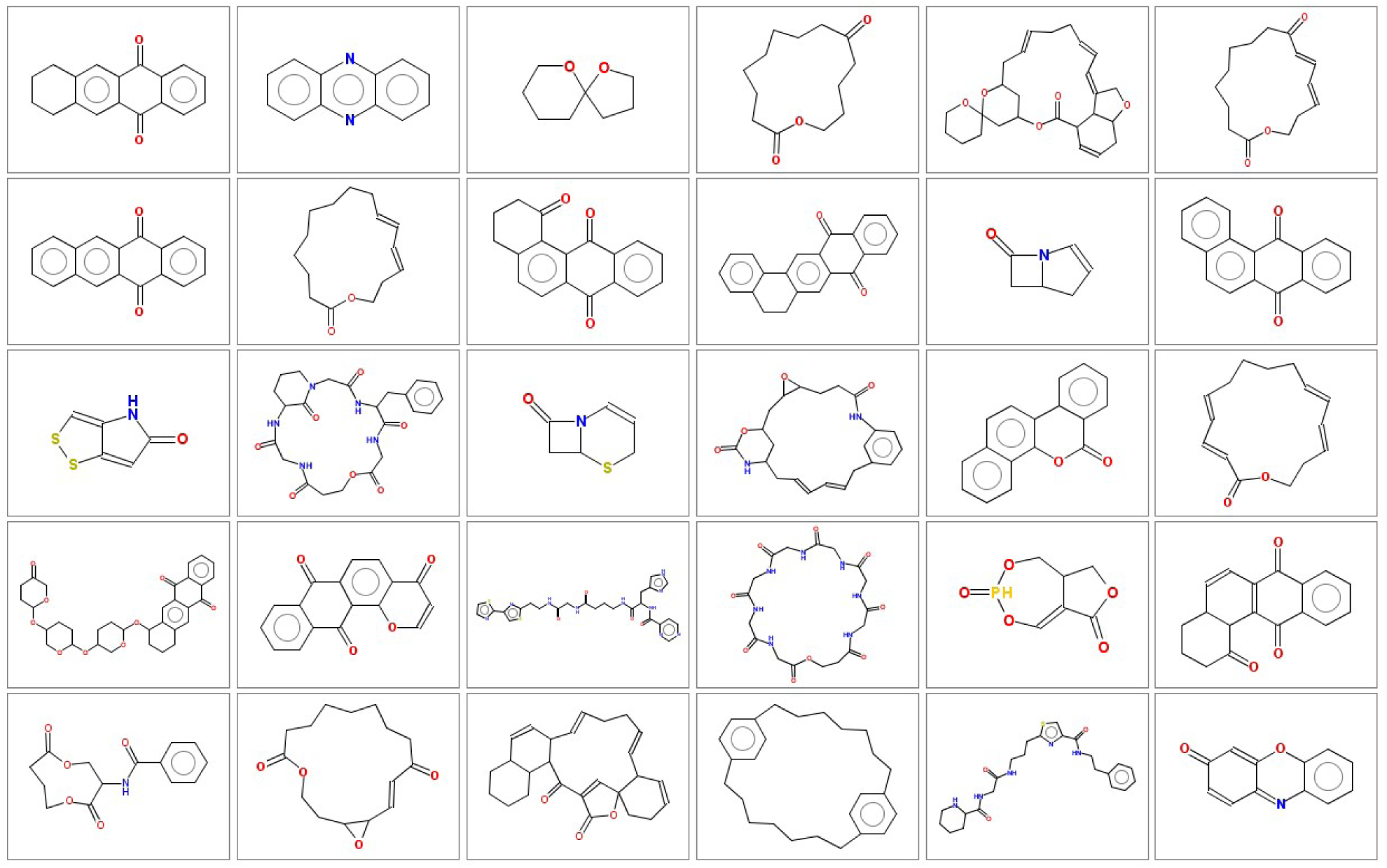
Typical scaffolds from molecules produced by bacteria.

As discussed before the bacterial scaffolds significantly differ from the other three classes practically in all substructure features analysed. The unique features of bacterial metabolites were also described in our recent analysis of frequency of functional groups in NPs produced by different classes of organisms. (Ertl and Schuhmann, 2019) and can to some extent be explained by an obvious tendency of bacteria to produce polyketide structures at least when cultivated at laboratory conditions.

The conclusions obtained in this analysis may serve as a guide when selecting or designing scaffolds for synthesis of “non-natural” natural products (i.e. molecules not produced by living organisms, but containing the same structural features as NPs) and be useful as central cores for synthesis of focused NP-like libraries.

## References

Carroll, A. R. et al. (2019) ‘Marine natural products’, Natural product reports, 36(1), pp. 122–173.

Davison, E. K. and Brimble, M. A. (2019) ‘Natural product derived privileged scaffolds in drug discovery’, Current opinion in chemical biology, 52, pp. 1–8.

‘Dictionary of natural products 27.1, CRC Press, Taylor & Francis Group, Boca Raton, FL, USA’, (no date). Available at: https://www.crcpress.com/go/the_dictionary_of_natural_products.

Elshahawi, S. I. et al. (2015) ‘A comprehensive review of glycosylated bacterial natural products’, Chemical Society reviews, 44(21), pp. 7591–7697.

Ertl, P. (2014) ‘Intuitive ordering of scaffolds and scaffold similarity searching using scaffold keys’, Journal of chemical information and modeling, 54(6), pp. 1617–1622.

Ertl, P. and Schuffenhauer, A. (2008) ‘Cheminformatics analysis of natural products: Lessons from nature inspiring the design of new drugs’, in Petersen, F. and Amstutz, R. (eds) Natural Compounds as Drugs, Vol II. Birkhaeuser Verlag, pp. 217–235.

Ertl, P. and Schuhmann, T. (2019) ‘A Systematic Cheminformatics Analysis of Functional Groups Occurring in Natural Products’, Journal of natural products, 82(5), pp. 1258–1263.

Grabowski, K., Baringhaus, K.-H. and Schneider, G. (2008) ‘Scaffold diversity of natural products: inspiration for combinatorial library design’, Natural product reports, 25(5), pp. 892–904.

Harvey, A. L., Edrada-Ebel, R. and Quinn, R. J. (2015) ‘The re-emergence of natural products for drug discovery in the genomics era’, Nature Reviews Drug Discovery, pp. 111–129. doi: 10.1038/nrd4510.

Koch, M. A. et al. (2005) ‘Charting biologically relevant chemical space: a structural classification of natural products (SCONP)’, Proceedings of the National Academy of Sciences of the United States of America, 102(48), pp. 17272–17277.

Newman, D. J. and Cragg, G. M. (2016) ‘Natural Products as Sources of New Drugs from 1981 to 2014’, Journal of Natural Products, pp. 629–661. doi: 10.1021/acs.jnatprod.5b01055.

Rodrigues, T. et al. (2016) ‘Counting on natural products for drug design’, Nature chemistry, 8(6), pp. 531–541.

Sauer, W. H. B. and Schwarz, M. K. (2003) ‘Size Doesn’t Matter: Scaffold Diversity, Shape Diversity and Biological Activity of Combinatorial Libraries’, CHIMIA International Journal for Chemistry, pp. 276–283. doi: 10.2533/000942903777679253.

Sterling, T. and Irwin, J. J. (2015) ‘ZINC 15--Ligand Discovery for Everyone’, Journal of chemical information and modeling, 55(11), pp. 2324–2337.

Tan, D. S. (2005) ‘Diversity-oriented synthesis: exploring the intersections between chemistry and biology’, Nature chemical biology, 1(2), pp. 74–84. Taxonomy - NCBI (no date). Available at: https://www.ncbi.nlm.nih.gov/taxonomy.

Wetzel, S. et al. (2007) ‘Cheminformatic Analysis of Natural Products and their Chemical Space’, CHIMIA International Journal for Chemistry, pp. 355–360. doi: 10.2533/chimia.2007.355.

